# pyNeurode: a real-time neural signal processing framework

**DOI:** 10.1101/2022.01.18.476764

**Authors:** Wing-kin Tam, Matthew F. Nolan

## Abstract

Accurate decoding of neural signals often requires assigning extracellular waveforms acquired on the same electrode to their originating neurons, a process known as spike sorting. While many offline sorters are available, accurate online sorting of spikes with many channels is still a challenging problem. Existing online sorters either use simple algorithms with low accuracy, can only process a handful of channels, or depend on a complex runtime environment that is difficult to set up. We have developed a state-of-the-art online spike sorting platform in Python that enables large-scale, fully automatic real-time spike sorting and decoding on hundreds of channels. Our system is cross-platform and works seamlessly with the Open Ephys suite of open-source hardware and software widely used in many neuroscience laboratories worldwide. It also comes with a user-friendly graphical user interface to monitor the cluster quality, spike waveforms and neuronal firing rate. Our platform has comparable accuracy to offline sorters and can achieve an end-to-end sorting latency of around 160 ms for 128-channel signals. It will be useful for research in fundamental neuroscience, closed-loop feedback neuromodulation and brain-computer interfaces.

## I. INTRODUCTION

Neurons in the brain communicate with each other through action potentials. In extracellular recordings with electrodes, these action potentials appear as spikes in the signal traces. The timing and firing rates of these spikes convey important information about the internal states of the brain. Analyzing this kind of neural activity in real-time can lead to many practical applications. For example, it can be used in a closed-loop feedback protocol to test hypotheses about brain functions, to build brain-computer interfaces enabling tetraplegic patients to control their environment or to deliver clinical intervention based on a particular pattern of neural activity.

However, analyzing many neural signals in real-time is challenging and involves multiple steps [1]. Neural signals are typically acquired at a high sampling rate of 20-40kHz. The signals are then filtered in the spike frequency range of 300-6000Hz. A certain threshold is then set to detect the spike waveforms. Since one electrode may record the signals from multiple neurons, it is essential to assign the spike waveforms to their originating neurons through clustering, a process known as “spike sorting”. The sorted spike timings are then used to estimate the firing rates of neurons. With the arrival of high-density silicon probes such as the Neuropixel [2], [3], more neuroscience laboratories are making large-scale recordings with hundreds of channels. Analyzing these amounts of signals efficiently in real-time is a demanding engineering problem.

Current toolkits for real-time neural signal processing suffer from significant drawbacks. First, although many offline spike sorters exist [4], only a few work online. Those that are designed for online use often use simpler algorithms with limited sorting accuracy [5], [6]. Online spike sorting of many channels also needs to be fully automatic as human intervention is not practical. Second, many online sorters emphasize performance over usability and use a compiled language such as C/C++ (e.g. [7]). Setting up their compilation and running environment is often not easily accessible due to many external library dependencies. Third, many online sorting systems are difficult to extend and customize. Many exist as proprietary software tailored for a particular data acquisition hardware (e.g. SpikeSort 3D for Neuralyx system, OmniPlex for Plexon, Spike2 for CED hardware etc.). Others are created as a monolithic application with tight coupling between components (e.g. [8]), so it is difficult for end-users to add new functionalities. However, online spike sorting is only useful if it is integrated as a component of a larger application, e.g. for closed-loop stimulation. Extensibility is not just a good-to-have feature but should be a core functionality of an online spike sorter.

To overcome these limitations, we have created pyNeurode (Python + neuron + node), a real-time neural signal processing framework for efficient online sorting of large-scale recordings. It uses the ISO-SPLIT algorithm from Mountainsort [9], [10] and so does not require setting of any free parameters. Its accuracy is also comparable with many offline sorters. pyNeurode is implemented in Python and is cross-platform compatible. It adopts a multi-process architecture to utilize the many computing cores of modern CPUs fully. It can sort 128 channels of recordings with around 160 ms latency. Each analysis process communicates via a message queue so additional functionality can be easily added without knowing the implementation details of other components. It also comes with a graphical user interface (GUI) to visualize the waveforms and principle components of sorted neuronal clusters, and the firing rate of each sorted spike in real-time. pyNeurode is designed to be a general real-time neural processing framework to enable closed-loop feedback experiments.

## II. METHODOLOGY

Our framework is designed to be easy to use and extend, able to fully utilize the multi-core processing power of modern computers, and provide accurate spike sorting and decoding of a large number of channels of neural signals.

### A. System architecture

Our framework is implemented in Python, a popular programming language among neuroscientists. It is relatively easy to learn and has a large body of third party libraries for data analysis and machine learning. By implementing our framework in Python, we take advantage of the rich ecosystem Python offers and allow it to be easily integrated into existing data processing pipelines. Python as an interpreted language also does not need to be compiled beforehand. Setting up the compilation environment is one of the major hurdles for extending existing software. Our framework is also cross-platform. It can be run on Windows, Linux and macOS.

We adopt an actor-based system architecture to make our framework easy to use and fully utilize the available computing resources. In this architecture, actors are independent and have no explicit shared state. We thus avoid the problem of explicit synchronization. This design offers great flexibility because each actor can run at its own speed. An actor is called a processor in our framework. Each processor runs independently as a system process and manages its internal states, thereby overcoming the limitation imposed by Python’s global intepretator lock (GIL). The GIL restricts the Python interpreter to execute a single instruction at a time and prevents true parallelism. processors communicate with each other via message passing through inter-process queues. This design choice brings about multiple benefits. First, it obviates the need for explicit synchronization between processors so that they can run independently and concurrently at different speeds. Second, by sending discrete messages rather than sharing a monolithic object in memory, we uncouple the processors from each other and make extending the framework much more accessible. Third, using a message passing mechanism allows a processor to add additional types of data down the processing pipeline easily without changing the implementation of other processors, thus making the whole system more robust to changes.

An overview of the system architecture is shown in Fig 1. An existing data acquisition system supported by our framework is the Open Ephys ecosystem of boards and software [11]. Open Ephys provides an open-source, economic and research-grade acquisition board widely used by many neuroscience laboratories worldwide. The Open Ephys GUI is the companion software used with the Open Ephys Acquisition Board to acquire neural data. For achieving high spike sorting accuracy, the signals must be first whitened to remove correlated noises. We have implemented a new online whitening plugin for the Open Ephys GUI. We have also improved the existing dynamic spike detector plugin to be compatible with the whitened signals. Additional improvement has also been made to the ZeroMQ plugin of Open Ephys to relay the detected spike waveforms to our real-time signal processing framework. Within our framework, each processor has an internal queue. Messages are passed between processors and put into the destination queue until they are processed. Some of the messages can additionally be sent via a side chain to a special GUI process for visualization. Our application programming interface (API) is designed to be as straightforward as possible. Configuring and connecting different processors only needs a few lines of code. A source code listing showing the setup of a fully functional online spike sorting application is given in Fig. 2.

**Fig. 1.**
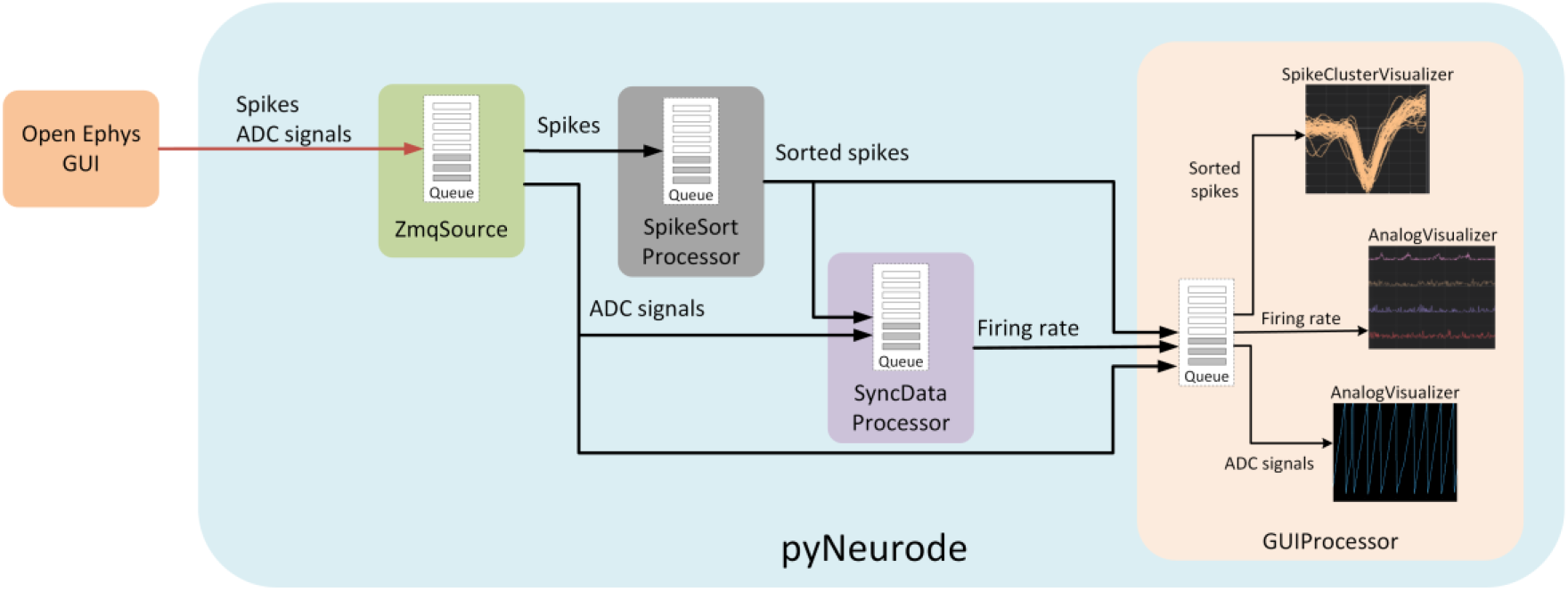
Overview of the system architecture. Spikes and analogue-to-digital converter (ADC) signals are streamed via ZeroMQ to the pyNeurode framework. Each processor in the framework runs in its own process and communicates via message passing through inter-process queues.

**Fig. 2.**
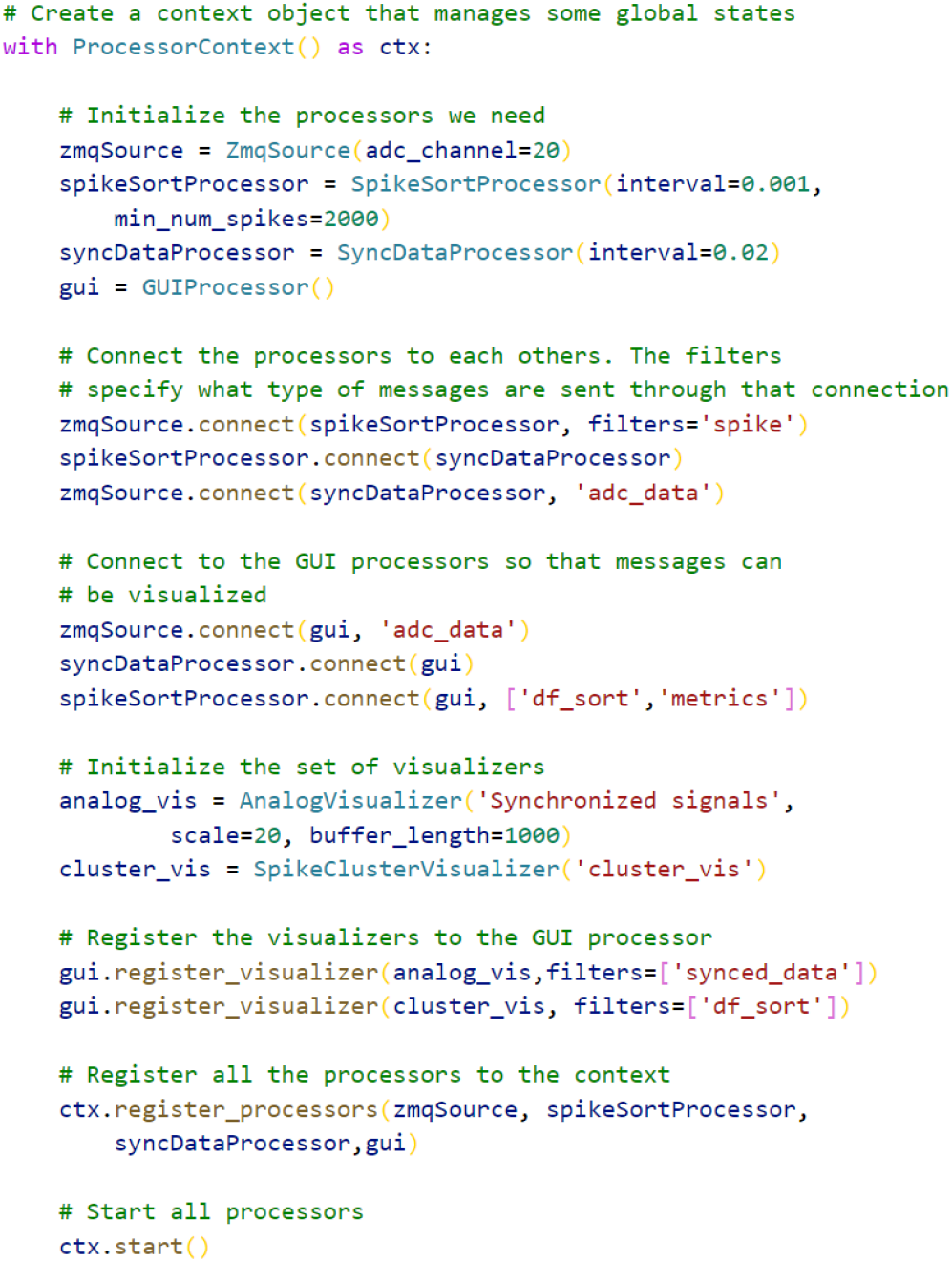
Source code of a fully functional online spike sorter, demonstrating the application programming interface and how the signal processors are connected. Comments explaining the functions of the code are preceded by a #.

### B. Online spike sorting

For brain-computer interface or closed-loop feedback control, it is vital that spike sorting be accurate and fully automatic. Human intervention is impractical for sorting a large number of channels in real-time. We adopted the ISO-SPLIT from MountainSort for online spike sorting [9]. ISO-SPLIT does not require the pre-specification of the number of clusters, but rather it relies on statistical tests to ensure each cluster is a unimodal distribution. In our framework, upon collecting an initial set of spike waveforms (the calibration set), the waveforms are first aligned to their negative peaks, then ISO-SPLIT is used to assign the spikes to different clusters. For each cluster, we calculate the average of the waveform and use it as a spike template. Subsequent spike waveforms are then matched to the existing template and assigned to the cluster with the highest correlation coefficient. Using template matching ensures we can sort a large number of spikes with minimal latency.

The sorted spike timings are then used to estimate each neuron’s firing rate and finally align with any behavioural variables (e.g. animal position). These aligned signals can then be fed into machine learning algorithms for decoding purposes.

### C. Graphical user interface

In many neuroscience experiments, it is necessary to monitor the states of the signal processing pipeline so that potential problems (e.g. inadvertent noise, connection issues, low task performance, poor quality sorting clusters) can be identified and corrected promptly. The analysis pipeline will also typically generate multiple types of intermediate signals at high speed. Therefore a neural signal processing framework must also provide an efficient way to visualize these many types of signals. It should also be easily extensible such that additional visualization can be added easily. To that end, we use Dear PyGUI [12], a new GUI framework that has extensive graphical acceleration built-in. It can visualize a large amount of information efficiently in real-time. In our framework, the GUI is nothing more than another processor. It spawns in its own process, receives messages via its input queue, and visualizes them according to their message types. Each type of message is associated with a certain visualizer object. The visualizer object is responsible for constructing the GUI window for that particular message type and updating the visuals when new data come in. The same message can be visualized differently by different visualizers, and multiple messages can be sent to the same visualizer for combined visualization. Each visualizer also works independently, allowing one to easily implement a new one by extending the base class and overriding two functions.

Currently, three visualizers that cover the essential functions are implemented. The Analog Visualizer allows displaying any multi-channel analogue signals. This is useful for showing the neural firing rate or other behavioural variables in an experiment. The Spike Cluster Visualizer shows the spike waveforms and the principal components of each sorted spike cluster. It allows the users to monitor the sorting accuracy. The Latency Visualizer displays processing latency for each of the processors. Additional visualizers can be easily implemented using the rich plotting function of the Dear PyGUI framework. A screenshot showing the GUI is shown in Fig. 3.

**Fig. 3.**
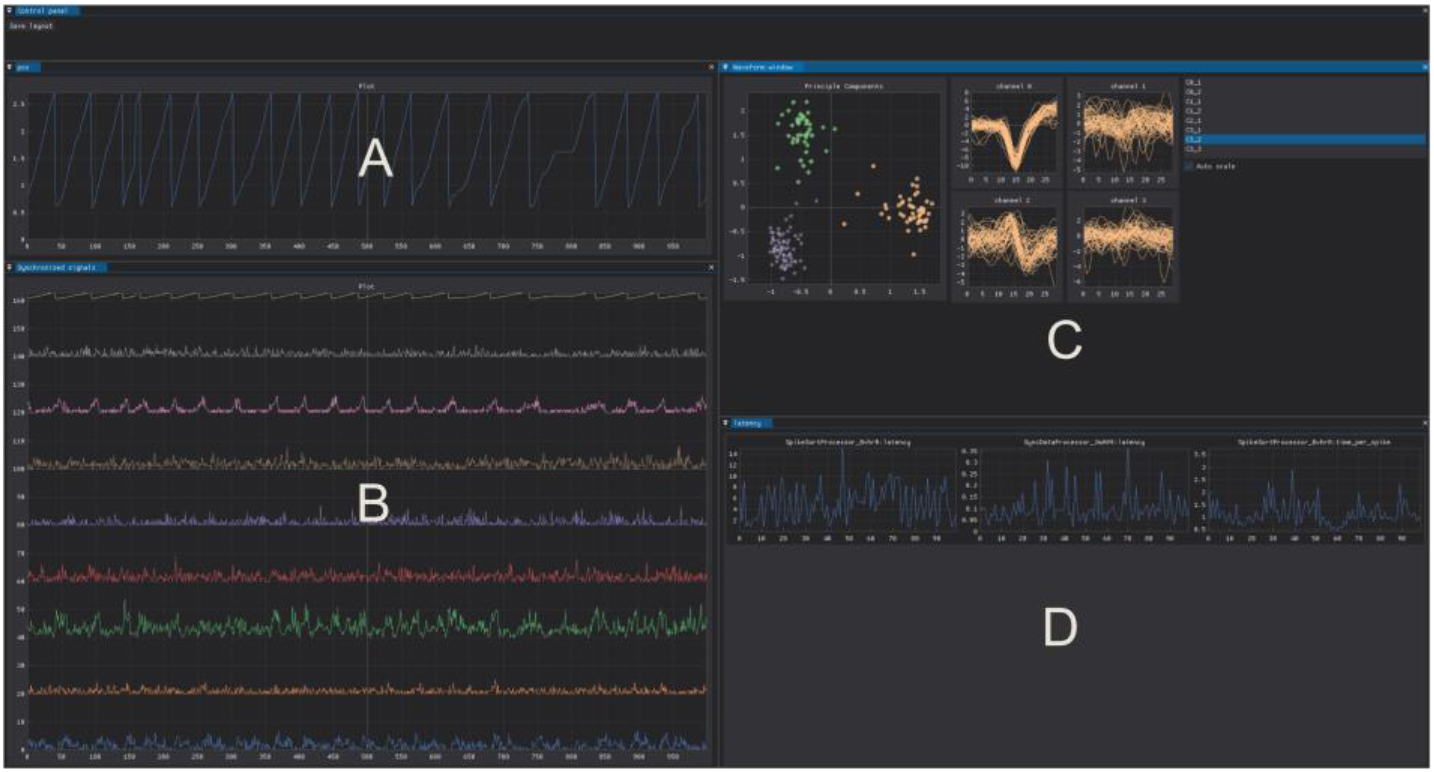
Graphical user interface (GUI) of the pyNeurode framework. (A) Animal position. (B) Firing rates of different neuronal clusters (C) Principal component plots (left) and waveform plots (right) of selected tetrode and one of its clusters, respectively. (D) Plots showing the latency of each processor.

### D. Evaluation procedures

#### 1) Spikes sorting accuracy

To evaluate the online spike sorting accuracy, we replayed a previously recorded 5 min 16-channel tetrode recording in the medial entorhinal cortex in Open Ephys [13], and let it pass through the entire data analysis pipeline, including filtering, whitening, spike detection in the Open Ephys GUI, and spike sorting in our framework. We then use the Spikeforest [4] package to sort the same recording with multiple offline sorters (Herding, Klusta, Ironclust and Spyking). Next, we compare the agreement scores of our online sorter and other offline sorters with Mountainsort. Detected spikes from different sorters are considered a match if they are within 0.4 ms of each other. The agreement score measures how many spikes are matched between two sorters.

#### 2) Latency

Latency is another important metric in real-time neural signal processing, especially for brain-computer interfaces or closed-loop feedback applications. Ideally, we want the processing to have as low latency as possible so that the subject’s intention is reflected quickly in the control or feedback. To measure the spike sorting latency, we measure the time difference between the moment a spike arrives in our framework and the moment that spike is sorted and its firing rate calculated. Hence it is the end-to-end latency in our framework and includes all internal communications overheads. Our GUI is also running and displaying the real-time neuronal firing rate and spike waveforms. During the latency measurement, pre-recorded data from [13] are replayed in Open Ephys GUI. Therefore, the latency measurements also take the computing load of a normal running Open Ephys GUI into account. The latency measurements are performed on a Windows 11 workstation PC with Intel Xeon E-2224G 4-core CPU and 47 GB of memory.

#### 3) Number of channels supported

With the arrival of large scale recording probes like the Neuropixels, recording many channels at once has become commonplace in many neuroscience laboratories. The multi-processing architecture of our framework is designed from the beginning to support large scale real-time spike sorting. We created large scale recordings by chopping our original 16-channel recording in time and then stacking them together as new channels. Hence the newly created channels will have realistic firing properties similar to real neurons. We measured the average latency of spike sorting in these synthetic recordings of different channel counts.

#### III. RESULTS AND DISCUSSION

The average agreement score between our online sorter and leading offline sorters is shown in Fig. 4A. Our agreement scores in some of the clusters (3, 20, 28) are even higher than that of the other offline sorters. One of the clusters (9) shows a significantly lower online agreement score. It is mainly because this cluster has a low firing rate, thus not enough of its waveforms are collected during the initial calibration phase training the spike templates. This problem can potentially be avoided by collecting more spikes for calibration. The cluster identification with our online sorter is comparable with those of the existing offline sorters.

**Fig. 4.**
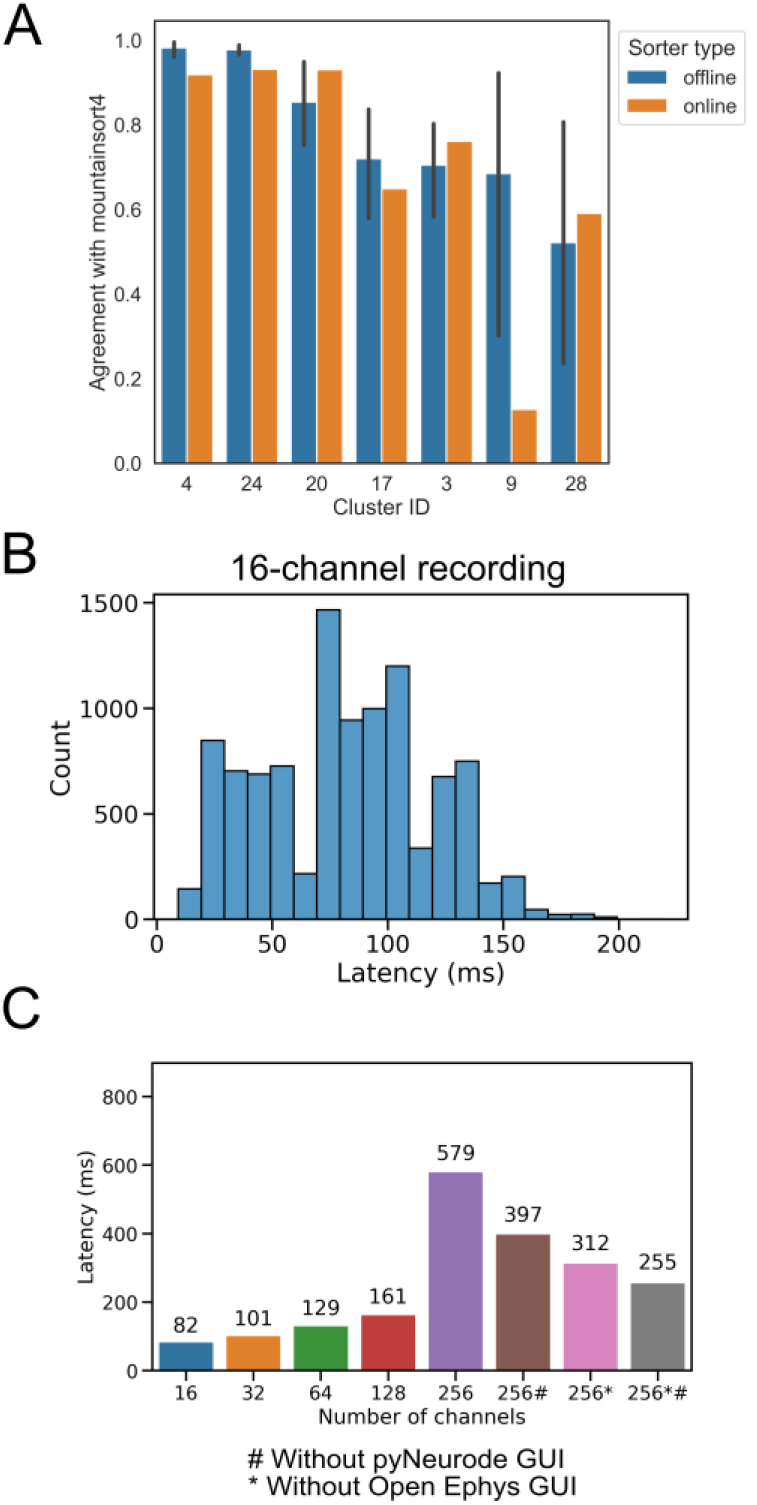
(A) The agreement score of our online sorter and the average score of four offline sorters (Herding, Klusta, Ironclust, Spyking) to Mountainsort. Cluster ID is the label of the cluster obtained by Mountainsort. The error bars show the 95% confidence interval of the mean. (B) Distribution of the sorting latency in a 16-channel recording for each spike. Note that many spikes are sorted at the same time and in a pipelined fashion. (C) Average spike sorting latencies in recordings with different numbers of channels. # means the pyNeurode GUI was disabled during measurement. * means the Open Ephys GUI was not used, but instead, previously recorded data packets were loaded from a file directly.

The end-to-end spike sorting latency within our framework during a 16-channel recording is shown in Fig. 4B. We can see that most of the spikes are sorted within 150 ms of arrival. For neural closed-loop control, latency in the order of a few hundred millisecond is generally considered to be acceptable [14], [15]. The average sorting latency in recordings with different numbers of channels is shown in Fig. 4C. Below 128 channels, the latencies increase with channel count. However, the increase in latency is much slower than the number of channels. For example, from 16 to 32 channels, the channel count is doubled, but the latency only increased by around 23%. This result demonstrates the strength of our parallel processing architecture. There is a large jump in the latency for 256 channels. The main reason for this is that the CPU is at its full load, thus slowing down the background thread that pulls the message items off the data queue. This limitation is mainly caused by our test machine’s low number of CPU processing cores (a mid-range Xeon E-2224G with 4 cores). If we disable the GUI or simulate the same rate of data in-flow by reading from a file instead of using the Open Ephys GUI, the latency can be reduced significantly, as seen in Fig. 4C. Both the GUI and the Open Ephys program run in a separate process and can be easily offloaded to additional CPU cores. Therefore, it is expected that by using higher-core CPUs, the latency for 256 channels can be further reduced.

In the future, we will continue to improve the performance and usability of our framework. We plan to rewrite some of the most computationally intensive parts of the code with Cython to improve its performance. Cython is a static compile for Python and can compile a subset of the Python language into C [14], thus improving its performance while maintaining a Python interface. We will also explore using a graphical processing unit to accelerate part of the pipeline [16]. We are also developing a node editor interface that allows end-users to easily drag and drop connections of different processors. This new interface will make the framework more user friendly for non-programmers. The source code of pyNeurode will be made freely available on Github after thorough testing and documentation.

## IV. ACKNOWLEDGEMENTS

This study is supported by the Simons Initiative for the Developing Brain.

## Notes

### Competing Interest Statement

The authors have declared no competing interest.

## REFERENCES

[1] H. G. Rey, C. Pedreira, and R. Quian Quiroga, “Past, present and future of spike sorting techniques,” Brain Res. Bull., vol. 119, pp. 106–117, 2015, doi: 10.1016/j.brainresbull.2015.04.007.

[2] J. J. Jun et al., “Fully integrated silicon probes for high-density recording of neural activity,” Nature, vol. 551, no. 7679, pp. 232–236, Nov. 2017, doi: 10.1038/nature24636.

[3] N. A. Steinmetz et al., “Neuropixels 2.0: A miniaturized high-density probe for stable, long-term brain recordings,” Science, vol. 372, no. 6539, Apr. 2021, doi: 10.1126/science.abf4588.

[4] J. Magland et al., “SpikeForest, reproducible web-facing ground-truth validation of automated neural spike sorters,” eLife, vol. 9, p. e55167, May 2020, doi: 10.7554/eLife.55167.

[5] G. Santhanam, M. Sahani, S. Ryu, and K. Shenoy, “An extensible infrastructure for fully automated spike sorting during online experiments,” Conf. Proc. Annu. Int. Conf. IEEE Eng. Med. Biol. Soc. IEEE Eng. Med. Biol. Soc. Annu. Conf., vol. 2004, pp. 4380–4384, 2004, doi: 10.1109/IEMBS.2004.1404219.

[6] S. Luan et al., “Compact standalone platform for neural recording with real-time spike sorting and data logging,” J. Neural Eng., vol. 15, no. 4, p. 046014, May 2018, doi: 10.1088/1741-2552/aabc23.

[7] S. Knieling et al., “An Unsupervised Online Spike-Sorting Framework.,” Int. J. Neural Syst., vol. 26, no. 0, p. 1550042, 2016, doi: 10.1142/S0129065715500422.

[8] T. K. T. Nguyen et al., “Closed-loop optical neural stimulation based on a 32-channel low-noise recording system with online spike sorting.,” J. Neural Eng., vol. 11, no. 4, p. 046005, Aug. 2014, doi: 10.1088/1741-2560/11/4/046005.

[9] J. F. Magland and A. H. Barnett, “Unimodal clustering using isotonic regression: ISO-SPLIT,” ArXiv150804841 Stat, May 2016, Accessed: Nov. 18, 2021. [Online]. Available: http://arxiv.org/abs/1508.04841

[10] J. E. Chung et al., “A Fully Automated Approach to Spike Sorting,” Neuron, vol. 95, no. 6, pp. 1381–1394.e6, 2017, doi: 10.1016/j.neuron.2017.08.030.

[11] J. H. Siegle, A. C. López, Y. A. Patel, K. Abramov, S. Ohayon, and J. Voigts, “Open Ephys: an open-source, plugin-based platform for multichannel electrophysiology,” J. Neural Eng., vol. 14, no. 4, p. 045003, 2017, doi: 10.1088/1741-2552/aa5eea.

[12] Dear PyGUI. [Online]. Available: https://github.com/hoffstadt/DearPyGui

[13] S. A. Tennant et al., “Analogue representation of a spatial memory by ramp-like neural activity in retrohippocampal cortex,” bioRxiv, p. 2021.03.15.435518, Mar. 2021, doi: 10.1101/2021.03.15.435518.

[14] R. Xu, N. Jiang, C. Lin, N. Mrachacz-Kersting, K. Dremstrup, and D. Farina, “Enhanced Low-Latency Detection of Motor Intention From EEG for Closed-Loop Brain-Computer Interface Applications,” IEEE Trans. Biomed. Eng., vol. 61, no. 2, pp. 288–296, Feb. 2014, doi: 10.1109/TBME.2013.2294203.

[15] R. T. Lauer, P. H. Peckham, K. L. Kilgore, and W. J. Heetderks, “Applications of cortical signals to neuroprosthetic control: a critical review,” IEEE Trans. Rehabil. Eng., vol. 8, no. 2, pp. 205–208, Jun. 2000, doi: 10.1109/86.847817.

[16] W. Tam and Z. Yang, “Neural Parallel Engine: A toolbox for massively parallel neural signal processing,” J. Neurosci. Methods, vol. 301, pp. 18–33, May 2018, doi: 10.1016/j.jneumeth.2018.03.004.

